# Perinatal Opioid Exposure Leads to Decreased Social Play in Adolescent Male and Female Rats: Potential Role of Oxytocin Signaling in Brain Regions Associated with Social Reward

**DOI:** 10.1101/2023.03.10.532122

**Authors:** Hannah J. Harder, Christopher T. Searles, Meghan E. Vogt, Anne Z. Murphy

## Abstract

Over the last two decades, the number of infants exposed to opioids *in utero* has quadrupled in the United States, with some states reporting rates as high as 55 infants per 1000 births. Clinical studies report that children previously exposed to opioids during gestation show significant deficits in social behavior, including an inability to form friendships or other social relationships. To date, the neural mechanisms whereby developmental opioid exposure disrupts social behavior remain unknown. Using a novel paradigm of perinatal opioid administration, we tested the hypothesis that chronic opioid exposure during critical developmental periods would disrupt juvenile play. As oxytocin is a major regulator of sociability, the impact of perinatal morphine exposure on oxytocin peptide and receptor expression was also examined. Juvenile play was assessed in vehicle- or morphine-exposed male and female rats at P25, P35, and P45. Classical features of juvenile play were measured, including time spent engaged in social play, time not in contact, number of pins, and number of nape attacks. We report that morphine-exposed females spend less time engaged in play behavior than control males and females, with a corresponding increase in time spent alone. Morphine-exposed females also initiated fewer pins and nape attacks. Oxytocin receptor binding was reduced in morphine-exposed females in the nucleus accumbens, a brain region critical for social reward. Together, these data suggest that females exposed to morphine during critical developmental periods are less motivated to participate in social play, potentially due to alterations in oxytocin-mediated reward signaling.

## 1. Introduction

The dramatic rise in maternal opioid use disorder in the United States over the last two decades has led to significant increases in the number of infants born with neonatal opioid withdrawal syndrome (NOWS) (Hirai et al., 2021). NOWS infants experience more complicated births and extended stays in the hospital, typically 9 days longer than non-exposed infants (Hirai et al., 2021). Although limited data are available regarding the impact of perinatal opioid exposure on behavioral outcomes, clinical studies have identified several cognitive, motor, and sensory deficits. Children with a history of NOWS have significantly lower cognitive and motor performance scores in early childhood (Baldacchino et al., 2014; Hunt et al., 2008) and are more likely to be diagnosed with learning disabilities than age- and demographically-matched controls (Fill et al., 2018; Maguire et al., 2016). Clinical literature further suggests *in utero* opioid exposure leads to long-lasting deficits in sociability. More specifically, children who were exposed to opioids prenatally score higher on assessments for autism spectrum disorder (ASD) and attention-deficit/hyperactivity disorder (ADHD) (Sandtorv et al., 2018). These observed deficits are especially prevalent in the “social difficulties” subscale, which assesses prosocial behavior, including the ability to form friendships, and social communication (Sandtorv et al., 2018). Similar studies investigating the impact of *in utero* opioid exposure also identified lower levels of social maturity during early childhood (Hunt et al., 2008).

Juvenile play is observed in most mammalian species and is essential for normal social development (see Bredewold and Veenema, 2018 or Veenema, 2012 for review). Participating in social play during adolescence facilitates the development of behavioral patterns that provide the framework for response to future challenges (Achterberg et al., 2019; Hol et al., 1996; Manduca et al., 2014). Indeed, isolation of rats during the juvenile period leads to deficits in adult social behavior, including decreased social motivation (Van Den Berg et al., 1999a), anogenital sniffing (Van Den Berg et al., 1999b), and social approach (Van Den Berg et al., 1999b; Van Den Berg et al., 2000). Therefore, social play is considered an indicator of general sociability, with adolescence being a critical period for its development.

Neuropharmacological studies utilizing acute opioid exposure in adulthood support a clear role for opioids in social attachment: systemic administration of μ-opioid receptor agonists enhances social play in male and female rats (Manduca et al., 2014; Vanderschuren et al., 1995a; Vanderschuren et al., 1995b; Vanderschuren et al., 1995c), and this effect is blocked by co-administration of μ-opioid receptor antagonists (Beatty and Costello, 1982; Jalowiec et al., 1989; Normansell and Panksepp, 1990; Siegel and Jensen, 1986; Siegel et al., 1985; Trezza et al., 2011). Similarly, administration of the μ-opioid receptor-selective antagonist CTAP into the caudate putamen of female prairie voles completely inhibits the formation of a partner preference (Burkett et al., 2011). In contrast, preclinical studies investigating the impact of perinatal opioid treatment on social play are limited and often contradictory, with reports of increased (Buisman-Pijlman et al., 2009; Hol et al., 1996; Niesink et al., 1999), decreased (Chen et al., 2015), and no effect (Minakova et al., 2022) on social behavior. These contradictions are likely due to differences in both the stage of gestation in which opioid administration is initiated and the duration of exposure. For example, in the above-mentioned studies, administration paradigms range from daily injections beginning on day 8 of pregnancy [E8] and continuing to parturition (Buisman-Pijlman et al., 2009) to exposure from E3-E20 (Chen et al., 2015); still others administer opioids from E0 until weaning at postnatal day 21 [P21] (Minakova et al, 2022). Importantly, these opioid administration paradigms fail to recapitulate the clinical profile of maternal opioid use, which typically begins in late adolescence, *before* pregnancy, and decreases in the final weeks before birth (Smith and Lipari, 2017). Thus, although clinical studies have linked gestational opioid exposure to deficits in sociability, there is a clear need for the development of more translational preclinical models to begin to address the underlying neural mechanisms.

The neuropeptide oxytocin (OT) has been implicated as a critical neuromodulator of social behavior (Bredewold et al., 2014; Rigney et al., 2022). For example, oxytocin signaling within the nucleus accumbens facilitates social reward in mice (Choe et al., 2022; Dölen et al., 2013), rats (Smith et al., 2017), and prairie voles (Johnson et al., 2017; Keebaugh et al., 2015; Keebaugh and Young, 2011), and OT+ Fos expression in the supraoptic nucleus (SON) is positively correlated with the percent of time juvenile male and female rats engaged in play (Reppucci et al., 2018). Critically, chronic opioid exposure in male rats leads to decreased oxytocin levels in the SON (Laorden et al., 1997) and blood (Van de Heijning et al., 1991) due to direct inhibition of OT synthesis by μ-opioid receptor agonists, including morphine (Li et al., 2001). Together, these studies suggest that developmental exposure to opioids may attenuate sociability by suppressing OT expression and/or signaling, however, to date, this has not been examined.

The present studies tested the hypothesis that perinatal morphine exposure leads to alterations in juvenile play via changes in OT peptide and receptor expression. Importantly, these studies are the first to use a clinically relevant rodent model in which morphine is administered prior to, during, and immediately following parturition to investigate opioid-induced sociability deficits.

## 2. Methods

### Experimental subjects

Male and female Sprague Dawley rats (approximately two months of age) were used to generate offspring perinatally exposed to morphine or sterile saline (vehicle) for controls (Charles River Laboratories, Boston, MA). After weaning on P21, all rats were housed in Optirat GenII ventilated cages (Animal Care Systems, Centennial, Colorado, USA) in same-sex pairs with corncob bedding. Food (Lab Diet 5001 or Lab Diet 5015 for breeding pairs, St. Louis, MO, USA) and water were provided ad libitum throughout the experiment, except during testing. All studies were approved by the Institutional Animal Care and Use Committee at Georgia State University and performed in compliance with the National Institutes of Health Guide for the Care and Use of Laboratory Animals. All efforts were made to reduce the number of rats used in these studies and minimize pain and suffering.

### Perinatal opioid exposure paradigm

Adult female Sprague Dawley rats (P60) were implanted with iPrecio SMP-200 microinfusion minipumps; morphine administration was initiated one week later. Pumps were programmed to deliver 10 mg/kg of morphine three times a day, with doses increasing weekly by 2 mg/kg until 16 mg/kg was reached (Figure 1). One week after morphine initiation, females were paired with sexually-experienced males for two weeks to induce pregnancy. Morphine exposure to the dams continued throughout gestation. At E18, approximately three days before parturition, pumps switched to 2x/day dosing, as initial studies using 3x/day dosing led to increased pup mortality following birth. Dams continued to receive morphine after parturition, such that pups received morphine through their mother’s milk. Beginning at P5, morphine dosage was decreased by 2 mg/kg daily until P7, when the dams received 0 mg/kg morphine (i.e., sterile saline). This protocol closely mirrors the clinical profile of infants exposed to opioids *in utero,* in which supplemental opioid dosing following birth is used to minimize withdrawal (Kocherlakota, 2014). Vehicle rats were treated in an identical manner, except pumps were filled with sterile saline.

**Figure 1.**
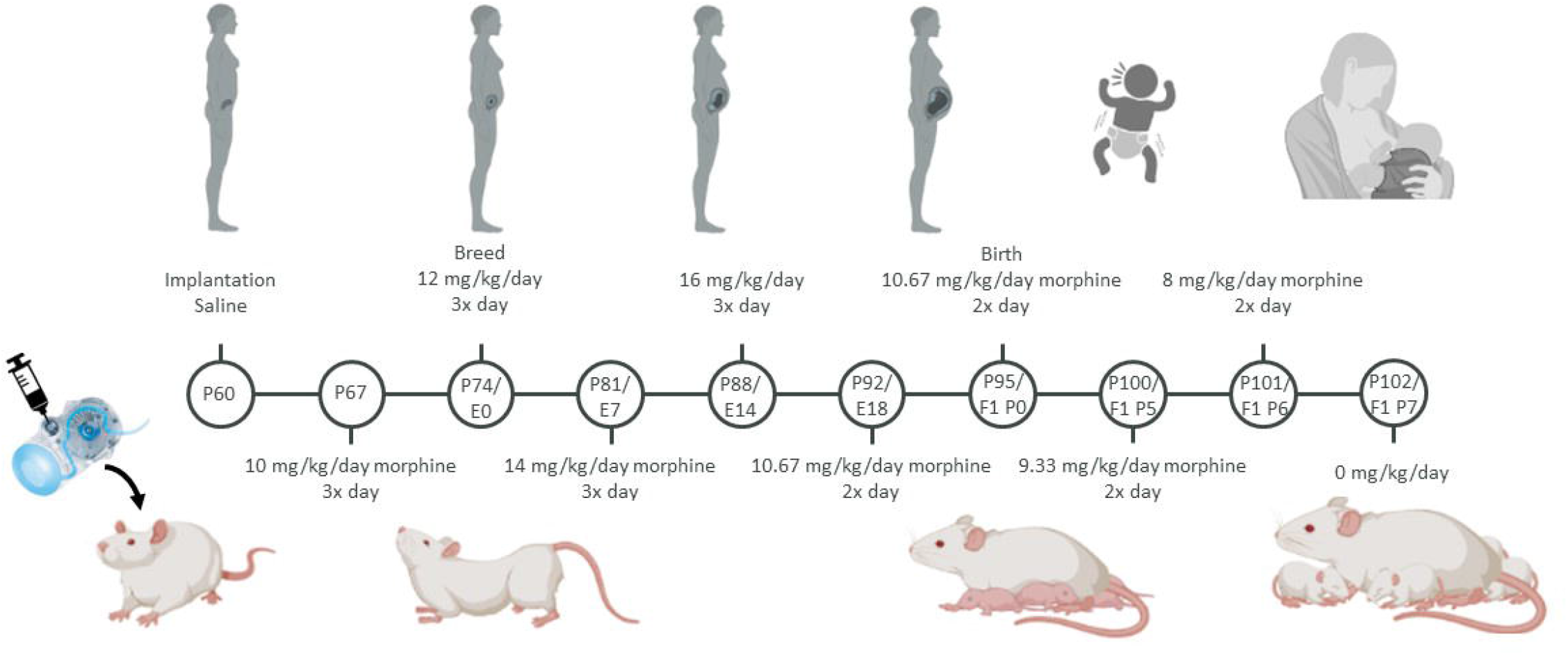
Schematic of the perinatal opioid exposure dosing paradigm. Created with BioRender.com
.

### Litter characteristics and dam/pup weight

Dams were weighed weekly during pregnancy to compare growth and weight gain between vehicle- and morphine-treated dams. At birth, litter size and sex balance were compared between vehicle- and morphine-exposed litters. At P4, anogenital distance (AGD) was measured using electronic calipers as a measure of potential endocrine disruption and/or differential androgen exposure (Schwartz et al., 2019). Pups were weighed every two days from P0 to P14, as neonatal morphine is associated with decreased growth and/or delayed developmental milestones (Najam and Panksepp, 1989).

### Maternal behavior

Maternal behavior was observed in the home cage during the light phase for one hour twice daily on P2, P4, and P6 to capture behavior during both high and low circulating levels of morphine. Following morphine cessation, maternal behavior was observed once daily on P8, P10, P12, and P14. Behaviors scored include nursing, licking/grooming, and time spent off pups.

### Social play paradigm

The typical developmental arc of juvenile social play is an inverted U-shape centered around P35 (Panksepp, 1981; Veenema et al., 2013); therefore, social play was measured three times over adolescence (P25, P35, and P45) to allow for the identification of any potential developmental shifts in morphine-treated offspring. Rats were separated from their cagemate for 24 hours before each trial to facilitate social play (Niesink and Van Ree, 1989; Vanderschuren et al., 1995d). Social behavior was videotaped for offline analysis for 12 minutes under red light. Social play was scored by two observers blind to condition. Five metrics were scored:

1. Time spent engaged in social play: total time spent chasing, pinning, and performing nape attacks.
2. Time spent in social interaction: total time spent sniffing or grooming their partner’s body.
3. Time spent not-in-contact: total time spent not interacting with each other (i.e., exploring the cage or walking around).
4. The number of pins: one rat lying on its dorsal surface with the other rat above it.
5. The number of nape attacks: one rat noses/rubs the neck of the other rat.

Behaviors 1-3 were scored per treatment- and sex-matched pair, while 4-5 were scored individually for each rat.

### Stress assessment

To assess separation-induced stress, rats were provided nestlet squares in their isolation cages. The nestlet square was pre-weighed, and the total percentage shredded was measured after twenty-four hours. Higher percentages of shredded material represent higher anxiety and/or compulsive behaviors resulting from isolation (Angoa-Perez et al., 2013). Previous studies have reported nestlet shredding is sensitive to anxiolytic drug administration, suggesting this is a valid indicator of anxiety-like behaviors in rodents (Li et al., 2006).

### Oxytocin immunohistochemistry

The effect of perinatal opioid exposure (POE) on oxytocin peptide expression was assessed using immunohistochemistry. Male and female morphine- and vehicle-exposed rats (P7, P14, or P30) were decapitated, brains rapidly removed, and drop-fixed in 4% paraformaldehyde for 24 hours, followed by 30% sucrose until sectioning. Fixed tissue was sectioned in a 1:6 series of 40-μm coronal sections with a Leica SM2010R microtome and stored in cryoprotectant at −20 °C. To visualize OT expression, free-floating sections were rinsed thoroughly in potassium phosphate buffer solution (KPBS), incubated in 3% hydrogen peroxide at room temperature for 30 minutes, and then rinsed in KPBS. Sections were then incubated in 1:10,000 mouse anti-oxytocin primary antibody (MAB5296, Millipore-Sigma) diluted in KPBS with 1%Triton-X overnight at room temperature. Following rinses in KPBS, sections were incubated in 1:600 biotinylated donkey anti-mouse secondary antibody (715-065-151, Jackson Immuno) diluted in KPBS with 0.4%Triton-X for one hour at room temperature. Following KPBS rinses, sections were incubated in an Avidin/Biotin solution (PK-6100, Vector Labs) for one hour at room temperature, followed by rinses in KPBS and sodium acetate. The sections were then incubated in a 3,3’-diaminobenzidine solution for 30 minutes, rinsed with sodium acetate and KPBS, and mounted onto slides. Slides were dehydrated using increasing concentrations of ethanol and cover-slipped. Oxytocin protein expression was quantified in the hypothalamic periventricular nucleus (PVN) and the supraoptic nucleus (SON), the main regions of oxytocin production and innervation. Four-six sections were analyzed, and the number of oxytocin neurons was counted and averaged per area for each rat.

Cresyl violet staining was performed on a subset of rats to investigate potential differences in overall cell number. Two-three 40-μm coronal sections per rat were mounted on slides and dipped in alternating solutions of 100% ethanol, xylene, 70% ethanol, 20% ethanol, water, cresyl violet/acetic acid solution, and differentiation solution (ethanol/acetic acid). Following cresyl violet staining, slides were cover-slipped and imaged for cell count analysis. Total cell counts were calculated using ImageJ across the hypothalamus by an experimenter blinded to condition.

### Oxytocin receptor autoradiography

To determine the impact of POE on oxytocin receptor binding, male and female morphine- and vehicle-exposed P7 rats were decapitated, brains were removed rapidly, hemisected, flash-frozen on dry ice, and stored at −80 °C. Frozen tissue was sectioned in a 1:6 series of 20-μm coronal sections at −20 °C with a Leica CM3050S cryostat. Sections were immediately mounted onto Superfrost Plus slides and stored at −80 °C until the time of the assay. To visualize OTR, sections were allowed to thaw to room temperature and fixed in 0.1% paraformaldehyde, followed by rinses in lx Trizma buffer. Slides were then placed in a tracer buffer containing 1-125-labeled OVTA ligand (Perkin Elmer NEX2540) for 60 minutes, followed by a series of rinses in 1x Trizma buffer containing MgCl_2_. After a final dip in cold dH_2_O, the tissue was allowed to dry at room temperature. Slides were then laid on cassettes with Kodak Biomax MR film for seven days. Films were developed using the Kodak X-Omat 2000A. Oxytocin-receptor binding was quantified from the film images using ImageJ software. [^14^C]-standards were used to convert uncalibrated optical density to disintegrations per minute (DPM). DPM was quantified in the nucleus accumbens (NAc), central amygdala (CeA), anterior cingulate cortex (ACC), and hippocampus due to their roles in social reward. Two slices per rat were analyzed, and the mean DPM per rat was calculated.

### Statistical analysis

Significant main effects of sex, treatment, age, and interaction effects were assessed using two- or three-way mixed models or repeated measures mixed models; p < 0.05 was considered significant. As repeated measures ANOVA cannot handle missing values, we analyzed our data by fitting mixed models with Greenhouse-Geisser correction (when the requirement of sphericity was not met) as implemented in GraphPad Prism 9.1.0 (Motulsky, 2021). Tukey’s or Sidak’s post hoc tests were conducted to determine significant mean differences between *a priori* specified groups. Due to the method of partitioning variance in linear mixed models, there is no universal method to calculate standardized effect sizes (e.g., η^2^ for ANOVA). Whenever possible, we report unstandardized effect sizes, which agree with recommendations for effect size reporting (Pek and Flora, 2018), including the guidance of the American Psychological Association Task Force on Statistical Inference (Wilkinson, 1999). Outlier points were defined as greater than two standard deviations from the mean of the treatment/sex/age group and were removed from future analysis. All experiments included both male and female offspring to investigate possible sex-specific effects of POE on social behavior. No differences were observed between rats of different litters in the same drug exposure group (i.e., morphine-exposed and vehicle-exposed); therefore, rats from multiple litters served as a single sample cohort.

## 3. Results

### Litter characteristics and dam/pup weight

To determine the effects of morphine exposure on developmental milestones, we investigated the weight gain of dams and pups, along with litter size, sex balance, and anogenital distance. All dams gained weight equally throughout pregnancy, regardless of treatment [F(5,33)=1.742, p=0.1525]. Vehicle-treated dams weighed, on average, 11.43 grams more than morphine-treated dams, approximately 5% difference in body weight (Figure 2A). At birth, litter size was significantly smaller for morphine-exposed dams [t(7)=2.38, p=0.0489] (Figure 2B). Vehicle-exposed litters consisted of, on average, 12 pups, while morphine-exposed litters consisted of an average of 9 pups (25.8% fewer). No effect of morphine on sex balance was observed [t(7)=0.48408, p=0.6453] (Figure 2C). On average, vehicle-exposed litters were comprised of 61.8% male pups, while morphine-exposed litters were 54.8% male.

**Figure 2.**
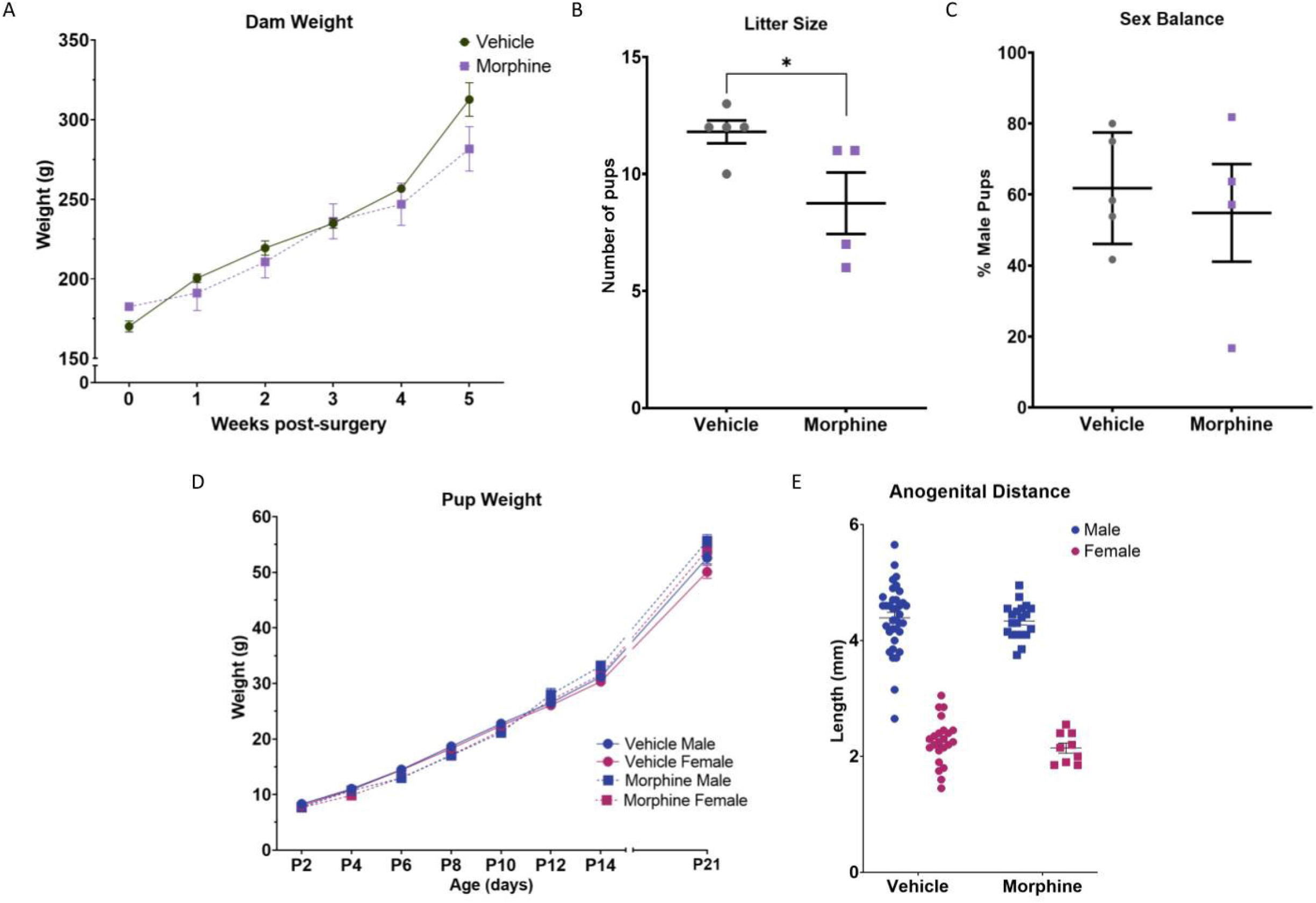
Perinatal opioid exposure does not lead to overt differences in litter characteristics or body weight. (A) Dams treated with morphine tend to weigh less later in pregnancy. (B) Litter size was significantly reduced in morphine-treated dams. (C) No difference in sex balance between vehicle- and morphine-treated dams. (D) Male and female offspring exposed to morphine weigh less than vehicle-exposed offspring from P2-P10 but weigh more than vehicle-exposed offspring at weaning. E. No effect of perinatal opioid exposure on anogenital distance. N_VEH M_ = 36-43, N_VEH F_ = 23-35, N_MOR M_ = 16-20, N_MOR F_ = 9-19. N_liters_: VEH, 5-7; MOR, 4. Graphs represent mean ± SEM. * = significant at p < 0.05.

There were no sex differences in pup weight between P2-P21; therefore, the sexes were collapsed. Analysis of pup weight identified a significant interaction between age and treatment [F(7,609)=6.443, p<0.001], with morphine-exposed pups weighing approximately 7% less than vehicle-exposed pups from P2-P10 but 6% heavier than vehicle-exposed pups at weaning (Figure 2D). Anogenital distance (AGD), measured on P4, was not significantly different between sex-matched vehicle- and morphine-exposed pups [F(1,84)=0.5853, p=0.4464] (Figure 2E). Average AGD for vehicle- and morphine-exposed males was 4.37 mm. Similarly, vehicle- and morphine-exposed females had an average AGD of 2.2 mm. Together, these results indicate no differences in developmental milestones between morphine- and vehicle-treated dams and their offspring.

### Maternal behavior

To examine the impact of morphine treatment on maternal behavior, we observed nursing and licking/grooming in the home cage of vehicle- and morphine-treated dams. Our analysis identified significant interactions of time and treatment for nursing [F(9,72)=3.551, p=0.0011] and licking and grooming [F(9,72)=2.648, p=0.0104]. Morphine-treated dams showed short-term decreases in nursing (30.1% less than vehicle-treated dams) when circulating levels of morphine were high (P2, P4, and P6 PM periods), but these differences were not observed at lower morphine levels (P2, P4, and P6 AM periods) or when morphine was no longer administered (after P8) (Figure 3A). Similar effects were observed with licking/grooming behavior (11.5% more than vehicle-treated dams) (Figure 3B). Overall, these data show that the effects of morphine on maternal behavior, both nursing and grooming, were minimal and acute.

**Figure 3.**
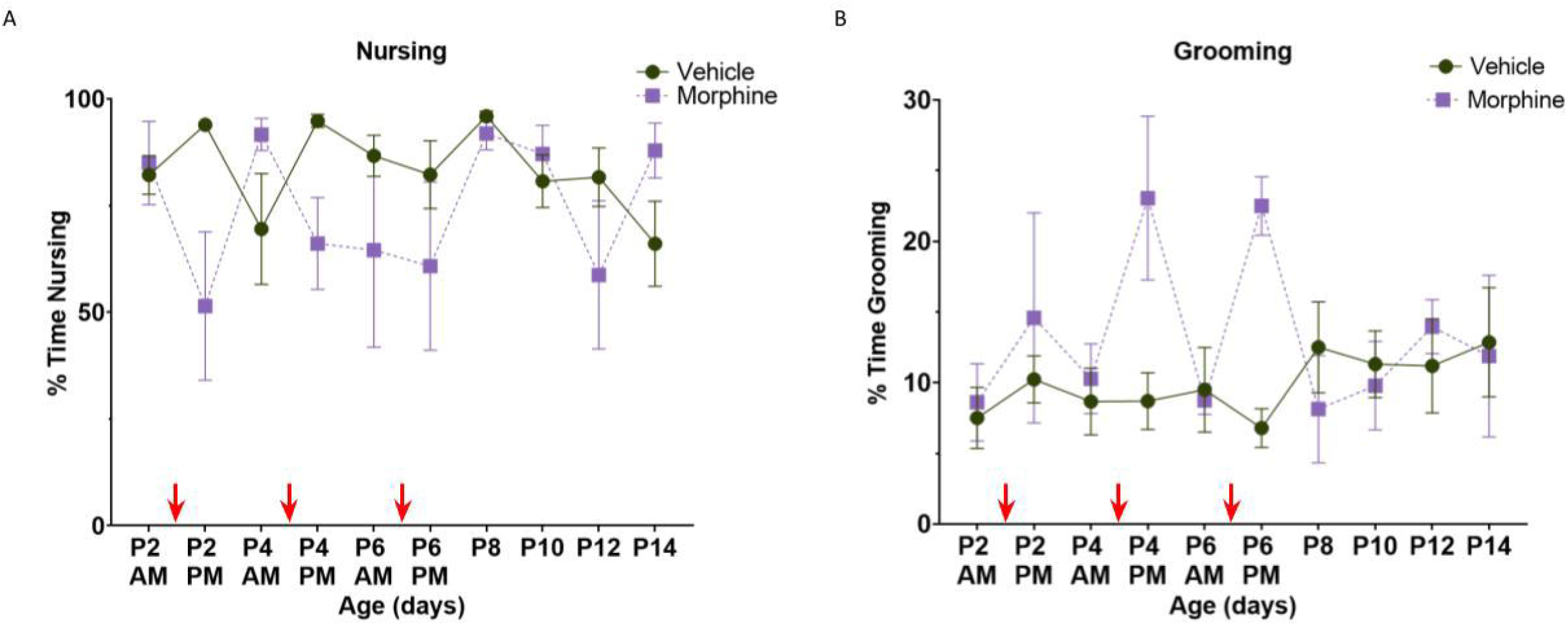
Maternal morphine treatment leads to short-term differences in maternal behavior. Dams treated with morphine spent less time nursing (A), and more time grooming (B) when morphine was administered (P2 PM, P4 PM, and P6 PM; indicated by red arrows). N_VEH_ = 7, N_MOR_ = 4. Graphs represent mean ± SEM. Red arrows indicate timing of morphine dose.

### Adolescent pup weights

To investigate the potential long-term effects of POE on body weight and its relationship with play behavior, we measured weight at P25, P35, and P45. There were no differences in weight between vehicle- and morphine-exposed offspring at any age [F(2,167)=2.904, p=0.0576] (Figure 4A), suggesting that POE does not lead to long-term differences in body weight or growth rate, as is sometimes observed with smaller litters (Parra-Vargas et al., 2020).

**Figure 4.**
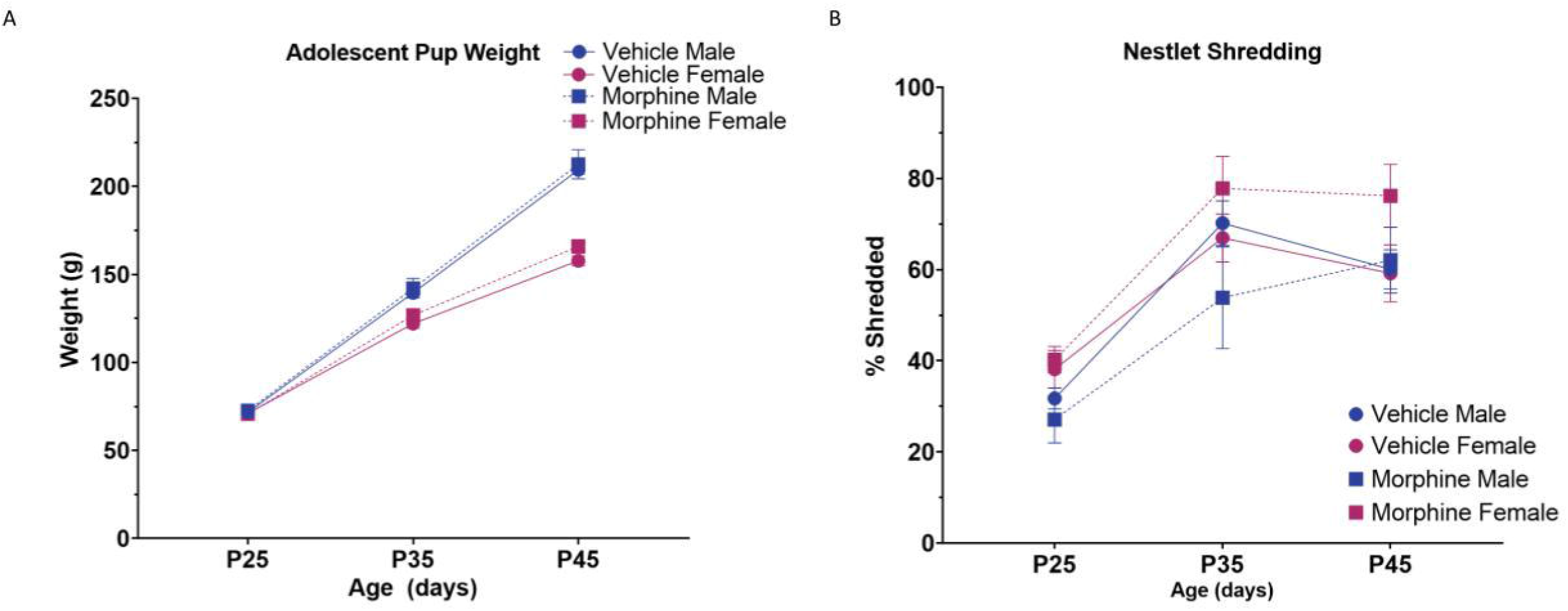
Adolescent weight **(A)** and nestlet shredding **(B)** do not differ between morphine- and vehicle-exposed male and female offspring. N_VEH M_ = 35, N_VEH F_ = 25, N_MOR M_ = 12, N_MOR F_ = 17. Graphs represent mean ± SEM.

### Nestlet shredding

Social isolation is a known stressor for rats; therefore, we used nestlet shredding to determine if repeated isolation prior to juvenile play was equally distressing for vehicle- and morphine-exposed offspring (Angoa-Perez et al., 2013). Higher percentages of shredded material are associated with higher anxiety and/or compulsive behaviors resulting from isolation (Angoa-Perez et al., 2013). Here, we found nestlet shredding increased as a function of repeated isolation [F(1.968,161.4)=86.24, p<0.0001]. This increase was comparable between vehicle- and morphine-exposed offspring (Figure 4B). Although not significantly different, morphine-exposed females shredded the most at all three timepoints, with an average percent shredded of 65%, 10-20% higher than all other groups. This suggests that morphine-exposed females may find repeated isolation more stressful than the other treatment-sex groups, potentially impacting their play behavior.

### Juvenile play and sociability

To determine the impact of perinatal morphine on juvenile social play, we next investigated five metrics of play: total time spent playing, time spent not-in-contact, time spent in social interaction, and the number of nape attacks and pins performed by each rat. Analysis of social play identified a three-way interaction of age, treatment, and sex [F(2,57)=3.299, p=0.0441]. At P35 and P45, morphine-exposed females had the lowest overall play times across all groups (Figure 5A). When averaged across all three developmental ages, morphine-exposed females spent the least time engaged in social play (Figure 5B). Specifically, morphine-exposed females played an average of 93.5 seconds in a 12-minute trial, 19.0% less than vehicle-exposed females. No sex difference in time spent not in contact was observed, so data were collapsed for analysis to increase power. ANOVA identified a significant effect of treatment [F(1,35)=4.164, p=0.0489] (Figure 5C). Morphine-exposed females spent the greatest amount of time not-in-contact across all three timepoints, with an average of 82.6 seconds, a 54.5% increase vs. vehicle-exposed females (Figure 5D). Morphine-exposed males had the second highest not-in-contact time, with an average of 57.6 seconds, or a 30.4% increase vs. vehicle-exposed males. To measure aspects of sociability independent from play, we assessed social exploration (licking, sniffing, and/or grooming the other rat). Analysis of social exploration identified a significant three-way interaction of age, treatment, and sex [F(2,57)=3.296, p=0.0442]. While social exploration time was approximately equal at P25 and P35, at P45, vehicle-exposed males spent 70.8 more seconds engaged in social exploration vs. morphine-exposed males, although this effect was not significant (Figure 5E). Females, regardless of treatment, showed comparable levels of social exploration at all three ages. When we analyzed average time spent across the three developmental timepoints, all groups showed comparable social exploration time, with a range of 526-550 seconds (Figure 5F). Together, these data indicate that morphine-exposed female rats are less engaged in social play, with a corresponding increase in not-in-contact time, and no corresponding alterations in social exploration.

**Figure 5.**
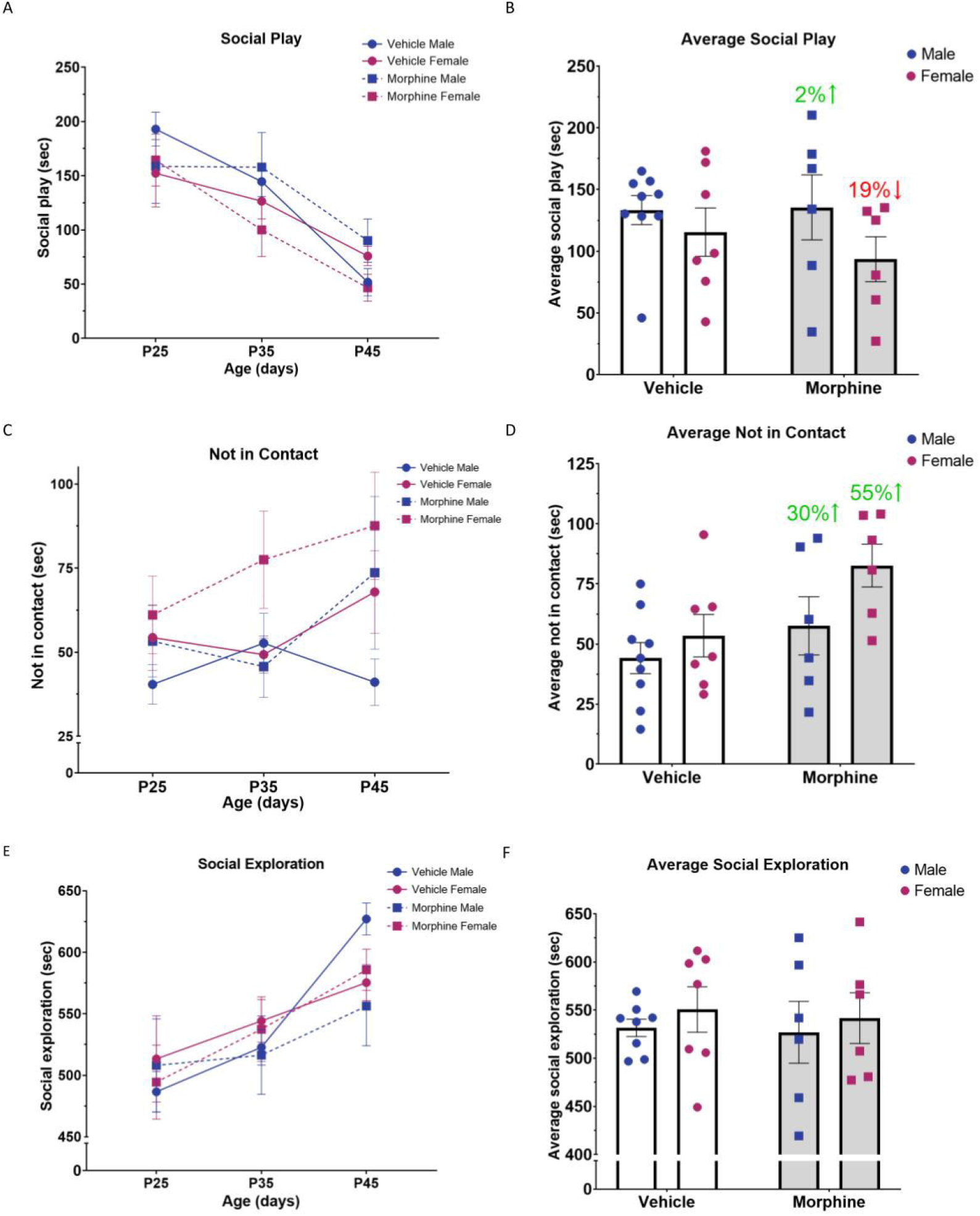
Morphine-exposed females show deficits in social behavior. **(A)** Morphine-exposed females play less than morphine-exposed males and vehicle-exposed male and females at P35 and P45 and **(B)** when averaged across all three ages. (C) Not-in-contact time was highest in morphine-exposed rats, regardless of sex and (D) when averaged across all three ages. (E) Vehicle-exposed males participate more in social exploration at P45, but there was no effect of treatment on average social exploration across all three ages **(F).** N_VEH M_ = 12 pairs, N_VEH F_ = 11 pairs, N_MOR M_ = 6 pairs, N_MOR F_ = 8 pairs. Graphs represent mean ± SEM. % represents change from sex-matched vehicle-exposed group.

### Nape attacks and pins

We next examined nape attacks and pins, two characteristic features of rat juvenile play. We began with nape attacks, which often serve to initiate a play bout and occur when a rat noses the other rat’s neck region. Analysis of nape attacks identified no treatment or sex differences; however, there was a significant effect of age [F(2,125)=17.30, p<0.0001], such that the number of nape attacks made at P35 and P45 was significantly lower than P25, regardless of treatment and sex group [P25 vs. P35, p=0.0007; P25 vs. P45, p<0.0001]. Morphine-exposed females had the lowest number of nape attacks at P35 and P45 (Figure 6A) and across all three ages, with an average of 10.4 nape attacks, a 28.6% decrease vs. vehicle-exposed females (Figure 6B). No difference in the average number of nape attacks was observed for vehicle- and morphine-exposed males.

**Figure 6.**
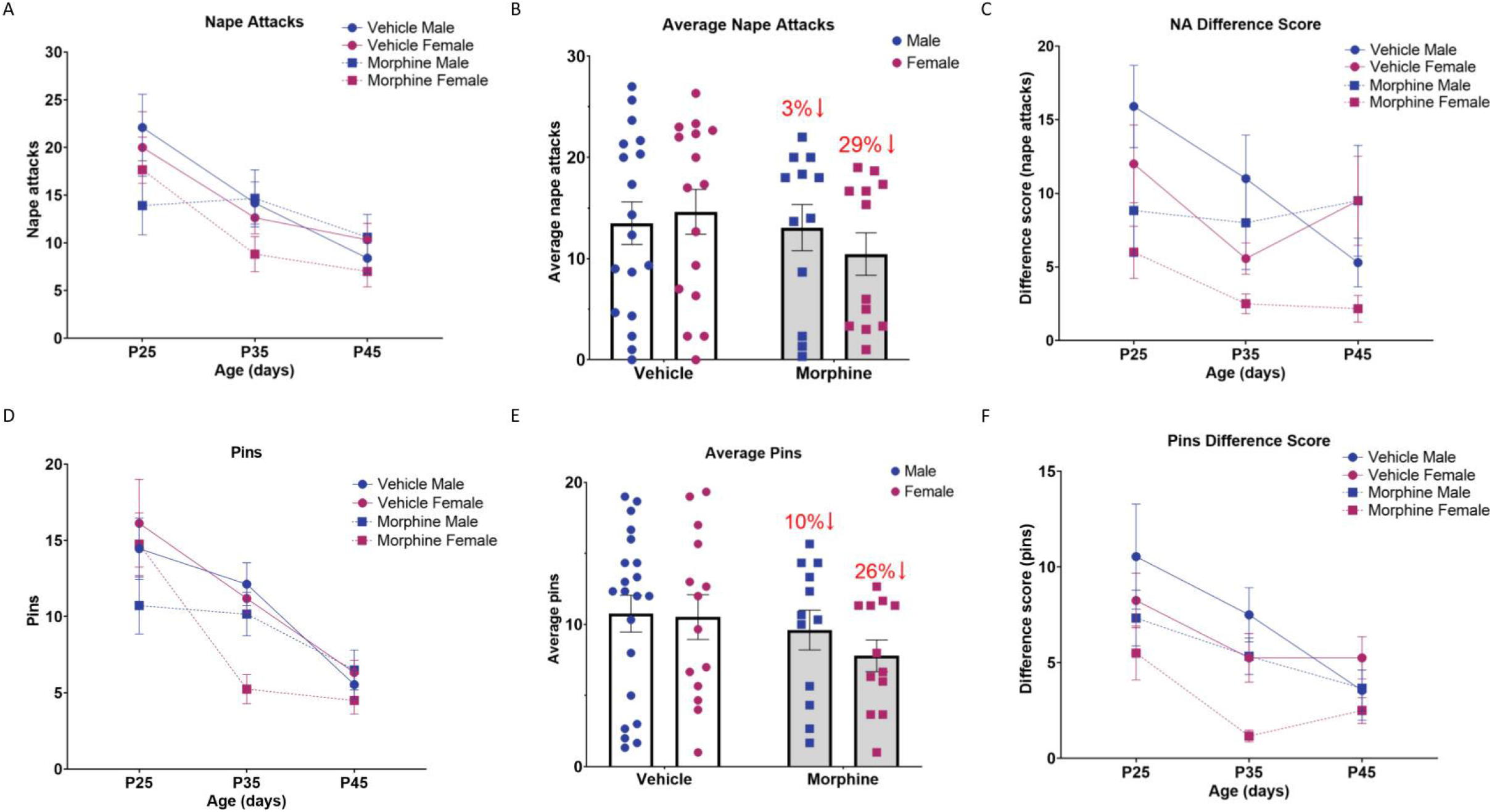
Morphine-exposed females display fewer characteristic features of social play. **(A)** Morphine-exposed females make the fewest nape attacks at P35 and P45 and (B) when averaged across all three ages. (C) Morphine-exposed males and females have significantly lower nape attack difference scores vs. vehicle-exposed males and females. (D) Morphine-exposed females make the fewest pins at P25, P35, and P45 and (E) when averaged across all three ages. (F) Morphine-exposed males and females have significantly lower pin difference scores vs. vehicle-exposed males and females, particularly morphine-exposed females. N_VEH M_ = 24, N_VEH F_ = 22, N_MOR M_ = 12, N_MOR F_ = 16. N_VEH M_ = 11 pairs, N_VEH F_ = 8 pairs, N_MOR M_ = 6 pairs, N_MOR F_ = 6 pairs. Graphs represent mean ± SEM. % represents change from sex-matched vehicle-exposed group.

To compare the number of nape attacks made by each member of treatment- and sex-matched pairs, we calculated a nape attack difference score: the absolute value of the difference in the number of nape attacks each rat made. Pairs with similar numbers of nape attacks have low difference scores, while disparate numbers create high difference scores. Asymmetry in the number of playful initiations is a hallmark of well-developed dominant-subordinate relationships between cagemates and is most common among adult male rats (Pellis et al., 1997). All treatment and sex groups had lower nape attack difference scores as a function of age [F(1.995,50.87)=2.129, p=0.0336], and morphine-exposed male and female rats had lower nape attack difference scores overall [F(1,27)=4.316, p=0.0474] (Figure 6C). This effect was particularly noticeable in morphine-exposed females, with an average difference score of 3.6, 60.6% lower than vehicle-exposed females. Morphine-exposed males showed a less extreme reduction in nape attack difference score, with an average score of 8.8, or 18.3% lower than vehicle-exposed males.

We next examined pins, which often serve to complete a play bout and occur when one rat is laying on its dorsal surface while the other rat hovers above. Analysis of pins identified a significant interaction of age and sex [F(2,124)=4.184, p=0.0175] as well as an overall effect of treatment [F(1,70)=4.647, p=0.0346] (Figure 6D). Morphine-exposed females made the fewest pins at both P35 and P45 and across all three ages, with an average of 7.8 pins, or a 25.8% decrease vs. vehicle-exposed females (Figure 6E). Morphine-exposed males made 10.7% fewer pins vs. vehicle-exposed males.

The difference score for pins was calculated similarly to the nape attack difference score, such that pairs of rats with similar numbers of pins have low difference scores, and pairs with disparate numbers of pins have high difference scores. Analysis of pins difference score identified significant effects of age [F(1.431, 37.92)=8.546, p=0.0024] and treatment [F(1,27]=4.493, p=0.0434]. Morphine-exposed rats, regardless of sex, had a lower pins difference score [VEH vs. MOR: p=0.0019]. This was especially noticeable in morphine-exposed females, who had the lowest difference score at P25, P35, and P45, 27.2% lower than vehicle-exposed females; morphine-exposed males also showed a 14.7% reduction in pins difference score vs. vehicle-exposed males (Figure 6F). Together, these data suggest that morphine-exposed rats, particularly females, display fewer characteristic features of juvenile play and that perinatal morphine disrupts the formation of normal cagemate relationships, as indicated by reduced dominant-subordinate hierarchies (see Table 1 for summary of results).

**Table 1.** Summary of behavioral results. Thick arrows represent significant comparisons of morphine- vs. vehicle-exposed rats.

|  | MOR M | MOR F |
| --- | --- | --- |
| Nestlet Shredding | — | ↑ |
| Social Play | — | ↓ |
| Not-in-Contact | ↑ | ↑ |
| Social Exploration | — | — |
| Nape Attacks | — | ↓ |
| Nape Attack Difference Score | ↓ | ↓ |
| Pins | ↓ | ↓ |
| Pins Difference Score | ↓ | ↓ |

### Oxytocin immunohistochemistry

Previous studies have strongly implicated oxytocin in regulating play and sociability. Therefore, we next examined the impact of perinatal opioid exposure on oxytocin expression using immunohistochemistry (Figure 7A). No significant sex differences were observed for the PVN or SON, so males and females were collapsed to increase power. Analysis of the total number of oxytocin-positive (OT+) cells in the PVN identified a significant interaction of age and treatment [F(2,53)=5.044, p=0.0099] (Figure 7B). Morphine-exposed rats had decreased OT+ cell count at P7, but an elevated OT+ cell count at P14 vs. vehicle-exposed rats. At P30, the number of OT+ cells were comparable between vehicle- and morphine-exposed rats (Table 2). Together, this suggests that perinatal morphine exposure alters OT+ cells in the PVN at P7 and P14, which is mostly normalized by P30.

**Figure 7.**
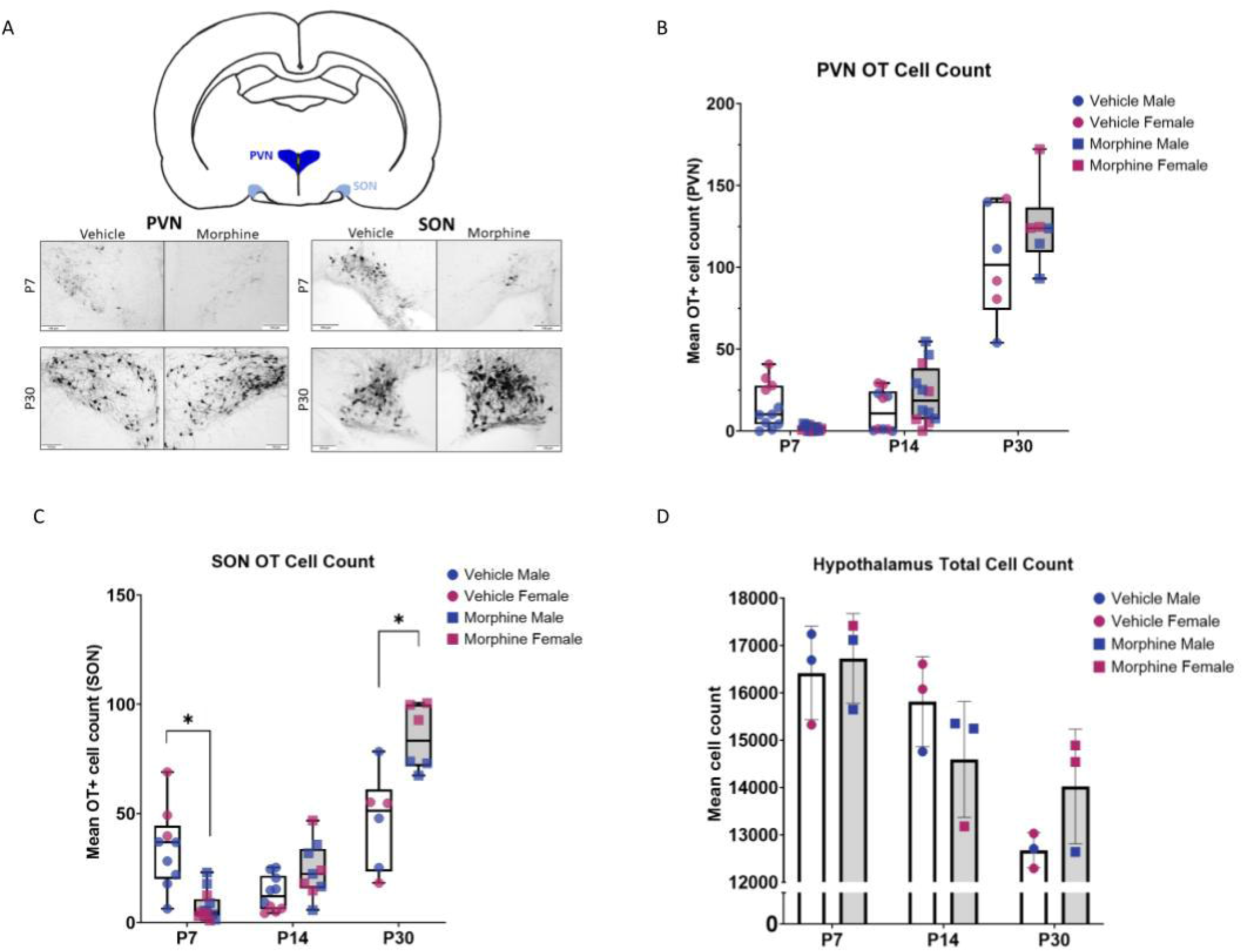
Morphine-exposed rats show decreased oxytocin expression in the PVN and SON. **(A)** Representative images of OT staining in the PVN and SON. (B) OT cell counts were reduced in morphine-exposed rats in the PVN at P7, increased at P14, and comparable at P30. (C) Similar results were observed in the SON at P7, P14, and P30. N_VEH_ = 6-11, N_MOR_ = 6-14. (D) Cresyl violet staining in the hypothalamus of a subgroup of rats showed no differences in overall cell count. N_VEH_ = 3, N_MOR_ = 3. Graphs (panels B-C) represent median ± IQR and (panel D) represent mean ± SEM. *: significant (p<0.05) Tukey’s post hoc analysis.

**Table 2.** 0T+ cell counts in the PVN for vehicle- and morphine-exposed rats at P7, P14, and P30. % represents percent change of morphine-exposed vs. vehicle-exposed rats.

| **PVN** | **P7** | **P14** | **P30** |
| --- | --- | --- | --- |
| **VEH** | 15.6 ± 13.8 | 12.6 ± 12.7 | 103.3 ± 34.6 |
| **MOR** | 1.6 ± 1.6<br>(↓ 89%) | 22.2 ± 17.9<br>(↑ 76%) | 125.4 ± 25.9<br>(↑ 21%) |

Similar results were observed in the SON. Analysis of the total number of oxytocin-positive cells identified a significant interaction of age and treatment [F(2,47)=23.70, p<0.0001] (Figure 7C). Again, morphine-exposed rats had decreased OT+ cell count at P7 and an elevated OT+ cell count at P14 vs. vehicle-exposed rats, which was maintained at P30 (Table 3).

**Table 3.** OT+ cell counts in the SON for vehicle- and morphine-exposed rats at P7, P14, and P30. *: significant (p<0.05) Tukey’s post hoc analysis. % represents percent change of morphine-exposed vs. vehicle-exposed rats.

| **SON** | **P7** | **P14** | **P30** |
| --- | --- | --- | --- |
| **VEH** | 34.0 ± 8.4 | 13.3 ± 8.0 | 46.6 ± 22.0 |
| **MOR** | 7.3 ± 6.6*<br>(↓ 79%) | 24.0 ± 12.4<br>(↑ 80%) | 84.6 ± 14.8*<br>(↑ 80%) |

Importantly, these differences in OT+ cell counts in the PVN and SON are not due to differences in overall cell number in the hypothalamus [F(2,12)=2.553, p=0.1192] (Figure 7D). Cresyl violet staining revealed minor differences in total cell count for vehicle- and morphine-exposed rats with no effect of sex at P7, P14, and P30. Together, this suggests that perinatal morphine exposure leads to initial decreases in oxytocin cell counts, with a potential compensatory increase in adolescence that is not due to overall differences in cell count in the hypothalamus.

### Oxytocin receptor autoradiography

The PVN and, to a lesser extent, the SON, provide oxytocinergic input to several regions implicated in social reward, including the central amygdala (CeA), anterior cingulate cortex (ACC), hippocampus (HPC), and nucleus accumbens (NAc). Given the significant reduction in oxytocin cell count at P7 in the PVN and SON, we next analyzed oxytocin receptor (OTR) binding using autoradiography in downstream regions involved in social reward. As observed for oxytocin cell count, there was no difference in receptor binding between males and females, so these groups were collapsed for the CeA, ACC, and HPC. No significant differences were observed between vehicle- and morphine-exposed rats at P7 in the CeA, ACC, or HPC (Figure 8A-C). However, analysis of OTR binding in the NAc identified a significant interaction of treatment and sex [F(1,9)=5.949, p=0.0374] (Figure 8D). Morphine-exposed females had the lowest receptor binding, with 91.7 DPM vs. 325.7 DPM for vehicle-exposed females, a 71.8% reduction (Figure 8E). No significant difference in OTR binding was noted for vehicle- and morphine-exposed males . Together with the oxytocin cell counts, these data suggest that perinatal morphine results in the dysregulation of oxytocin signaling and may contribute to the decreased rewarding nature of play observed in morphine-exposed female rats.

**Figure 8.**
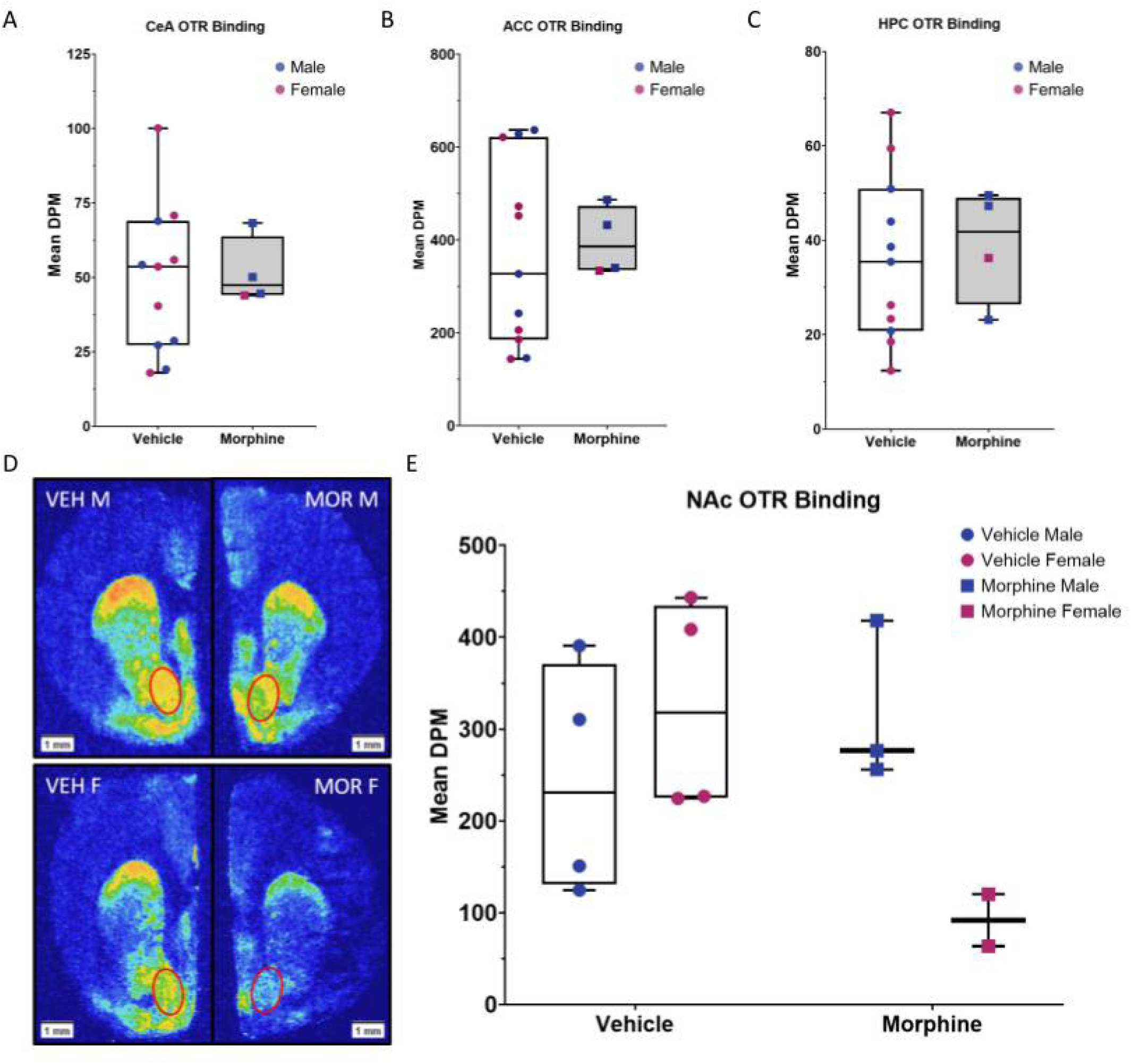
Morphine-exposed females show the lowest oxytocin receptor binding in the nucleus accumbens at P7. No significant differences in OTR binding were noted for the in the central amygdala **(A),** anterior cingulate **(B),** or hippocampus **(C).** N_VEH_ = 11, N_MOR_ = 4. **(D)** Representative images of the nucleus accumbens in VEH M, MOR M, VEH F, and MOR F. Area of quantification is circled. **(E)** Average DPM in the nucleus accumbens was lowest in morphine-exposed females. N_VEH M_ = 4, N_VEH F_ = 4, N_MOR M_ = 3, N_MOR F_ = 2. Graphs represent median ± IQR.

## 4. Discussion

The present study was designed to investigate the impact of perinatal morphine exposure on juvenile play behavior and oxytocin peptide and receptor expression. We hypothesized that exposure to opioids during fetal development would lead to decreased oxytocin signaling, in line with morphine’s known suppressive effects on oxytocin synthesis. Given the pro-social roles of oxytocin, we further hypothesized that morphine-induced changes in oxytocin signaling would negatively impact juvenile play. Our results showed a significant effect of treatment with reduced social play and altered oxytocin peptide and receptor expression that was most prevalent in females. More specifically, females perinatally exposed to morphine engaged in less social play, particularly as they aged, as indicated by decreased time spent in social play and increased not-in-contact time. Although morphine-exposed females also showed increased isolation-induced stress behavior, this is not likely to explain their decreased social play, as isolation is typically associated with increased play (Panksepp, 1981). Morphine-exposed females did not display concurrent social exploration deficits in agreement with previous studies reporting that the impact of acute μ-opioid receptor agonists during the juvenile period is limited to play and not social exploration (Manduca et al., 2014; Trezza et al., 2011; Van Ree and Niesink, 1983). Our perinatal morphine administration paradigm recapitulates the clinical profile of human infants exposed to opioids *in utero,* incorporating opioid exposure before and during gestation, and immediately after parturition. An essential part of this model is the inclusion of morphine before E15, the approximate date of μ-opioid receptor development in the fetal rat brain (Coyle and Pert, 1976). As there are potential trophic effects of opioids early in development (Kuhn et al., 1992), the use of a perinatal administration paradigm improves the clinical relevance and translatability of our findings to humans with *in utero* opioid exposure.

In the present study, lower OT+ cell counts in the PVN and SON were observed at P7 in morphine-exposed rats, likely due to morphine’s inhibitory effects on oxytocin synthesis. Morphine-exposed rats showed elevated OT cell count in the SON at P30, suggesting a potential “rebound” or compensatory increase. Mirroring these results, our autoradiography data revealed that morphine-exposed females have decreased oxytocin receptor binding at P7 in the nucleus accumbens, an important hub for reward signaling. Although not directly tested in the present study, we speculate that decreases in oxytocin expression and receptor binding observed in this study are likely due to the presence of morphine during critical periods of brain development and contributed to the observed deficits in adolescent social play. Opioids, particularly μ- and κ-receptor agonists, generally inhibit OT and vasopressin (AVP) neuronal firing, thereby suppressing peptide release (Douglas and Russell, 2001; Inenaga et al., 1990; Li et al., 2001; Lutz-Bucher and Koch, 1980; Van de Heijning et al., 1991), likely via presynaptic mechanisms (Clarke and Wright, 1984; Zhao et al., 1988). Interestingly, juvenile play also facilitates the development of inhibitory prefrontal cortex synapses necessary for cognitive and executive function (Bijlsma et al., 2022). This suggests that clinical reports of reduced social behavior and deficits in cognitive performance may be inter-related (Yeoh et al., 2019).

Our results are consistent with previous studies reporting that chronic opioid administration to adult male rats significantly decreases OT in both the PVN and SON (Laorden et al., 1997; Laorden et al., 1998; You et al., 2000). Together with the present results, this suggests that opioid-induced downregulation of oxytocin expression is not limited to specific developmental windows and is a likely mediator of opioid-induced alterations in sociability. Importantly, people with opioid use disorder have higher rates of social isolation, which often cyclically produces greater levels of drug use and further feelings of social isolation (Christie et al., 2021). This was particularly prevalent during the COVID-19 pandemic, in which rates of opioid use and overdose significantly increased (Alter and Yeager, 2020; Niles et al., 2021). Opioid-induced decreases in oxytocin production and binding may be one mechanism underlying this bidirectional, cyclic relationship between opioid exposure and social isolation. This connection may also suggest that children who were exposed to opioids *in utero* would be more susceptible to drug use and addiction later in life, as is seen in other preclinical models of gestational opioid exposure (Gagin et al., 1997; Ramsey et al., 1993; Shen et al., 2016; Torabi et al., 2017; Wu et al., 2009).

The results of the present study have implications for the clinical population of opioid-exposed infants that show sociability deficits, including social difficulties and inattention (Sandtorv et al., 2018). For example, children prenatally exposed to opioids were reported to have significantly higher scores on the “social difficulties” subscale of the Autism Spectrum Screening Questionnaire, including a lack of empathy and eye contact, no or few friends, and being poor at playing games (Sandtorv et al., 2018). These deficits are commonly observed in autism spectrum disorder and attention deficit hyperactivity disorder (Alessandri, 1992; Bredewold et al., 2014; Jordan, 2003; Reppucci et al., 2018), particularly in children. Alterations in OT signaling are implicated in social play deficits and ASD (Reppucci et al., 2018), including lower OT levels and polymorphisms in the *OXTR* gene (Meyer-Lindenberg et al., 2011; Zhang et al., 2017). OT administration has been reported to improve social functioning in children with ASD (Meyer-Lindenberg et al., 2011), suggesting the OT system is a prime therapeutic target for improving social play deficits in children born after *in utero* opioid exposure.

In summary, the current study provides evidence that perinatal morphine exposure disrupts early life oxytocin signaling in the social reward circuit, resulting in atypical adolescent social play. This effect was observed predominately in females. Specifically, we show that perinatal morphine decreases key features of social play and that this decrease is associated with a decrease in OT and OTR in regions implicated in social behavior and reward (see Figure 9 for a summary of proposed model). The use of a clinically relevant model of perinatal opioid exposure boosts the translational value of these studies in informing the development of potential clinical interventions for improving sociability in infants with *in utero* opioid exposure.

**Figure 9.**
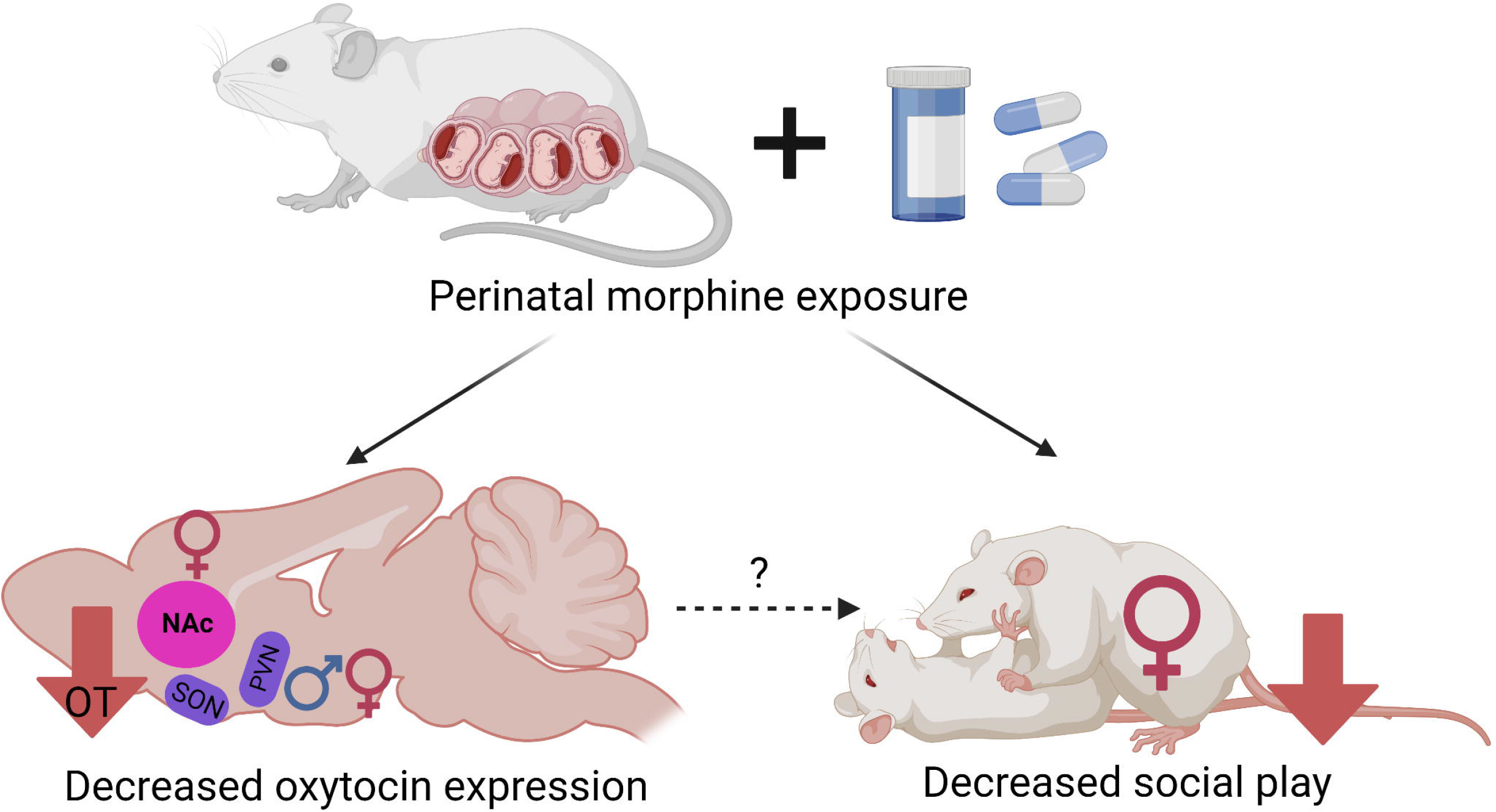
Model of proposed effect. Perinatal morphine exposure leads to decreased oxytocin expression in the PVN and SON in male and females, and decreased oxytocin binding in the NAc only in females. This suggests that perinatal morphine results in dysregulation of the oxytocin system during a critical period of social development, likely leading to long-term changes in juvenile play. Created with BioRender.com
.

## References

Achterberg, E.J.M., van Swieten, M.M.H., Houwing, D.J., Trezza, V., Vanderschuren, L.J.M.J., 2019. Opioid modulation of social play reward in juvenile rats. Neuropharmacology 159, 107332. https://doi.org/10.1016/i.neuropharm.2018.09.007.

Alessandri, S.M., 1992. Attention, play, and social behavior in ADHD preschoolers. J. Abnorm. Child Psychol. 20, 289–302. https://doi.org/10.1007/BF00916693.

Alter, A., Yeager, C., 2020. The consequences of COVID-19 on the overdose epidemic: Overdoses are increasing. Overdose Detection Mapping Application Program. https://odmap.org:4443/Content/docs/news/2020/ODMAP-Report-May-2020.pdf.

Angoa-Pérez, M., Kane, M.J., Briggs, D.I., Francescutti, D.M., Kuhn, D.M., 2013. Marble burying and nestlet shredding as tests of repetitive, compulsive-like behaviors in mice. J Vis Exp:50978. https://doi.org/10.3791/50978.

Baldacchino, A., Arbuckle, K., Petrie, D.J., McCowan, C., 2014. Neurobehavioral consequences of chronic intrauterine opioid exposure in infants and preschool children: a systematic review and meta-analysis. BMC Psychiatry 14, 104. https://doi.org/10.1186/1471-244X-14-104.

Beatty, W.W., Costello, K.B., 1982. Naloxone and play fighting in juvenile rats. Pharmacol. Biochem. Behav. 17, 905–907. https://doi.org/10.1016/0091-3057(82)90470-1.

Bijlsma, A., Omrani, A., Spoelder, M., Verharen, J.P.H., Bauer, L, Cornelis, C., de Zwart, B., van Dorland, R., Vanderschuren, L.J.M.J., Wierenga, C.J. 2022. Social play behavior is critical for the development of prefrontal inhibitory synapses and cognitive flexibility in rats. J Neurosci:JN-RM-0524-22. https://doi.org/10.1523/JNEUROSCI.0524-22.2022.

Bredewold, R., Smith, C.J.W., Dumais, K.M., Veenema, A.H., 2014. Sex-specific modulation of juvenile social play behavior by vasopressin and oxytocin depends on social context. Front. Behav. Neurosci. 8, 216. https://doi.org/10.3389/fnbeh.2014.00216.

Bredewold, R., Veenema, A.H., 2018. Sex differences in the regulation of social and anxiety-related behaviors: insights from vasopressin and oxytocin brain systems. Curr Opin Neurobiol 49, 132–140. https://doi.org/10.1016/i.conb.2018.02.011.

Buisman-Pijlman, F.T.A., Gerrits, M.A.F.M., Van Ree, J.M., 2009. Increased opioid release in specific brain areas in animals exposed to prenatal morphine and emotional stress later in life. Neuroscience 159, 405–413. https://doi.org/10.1016/j.neuroscience.2008.11.010.

Burkett, J.P., Spiegel, L.L., Inoue, K., Murphy, A.Z., Young, L.J., 2011. Activation of μ-opioid receptors in the dorsal striatum is necessary for adult social attachment in monogamous prairie voles. Neuropsychopharmacology 36, 2200–2210. https://doi.org/10.1038/npp.2011.117.

Chen, H.H., Chiang, Y.C., Yuan, Z.F., Kuo, C.C., Lai, M.D., Hung, T.W., Ho, I.K., Chen, S.T., 2015. Buprenorphine, methadone, and morphine treatment during pregnancy: Behavioral effects on the offspring in rats. Neuropsychiatr. Dis. Treat. 11, 609–618. https://doi.org/10.2147/NDT.S70585.

Choe, K.Y., Bethlehem, R.A.I., Safrin, M., Dong, H., Salman, E., Li, Y., Grinevich, V., Golshani, P., DeNardo, L.A., Peñagarikano, O., Harris, N.G., Geschwind, D.H., 2022. Oxytocin normalizes altered circuit connectivity for social rescue of the Cntnap2 knockout mouse. Neuron 110, 795–808.e6. https://doi.org/10.1016/j.neuron.2021.11.031.

Christie, N. C., 2021. The role of social isolation in opioid addiction. Social Cognitive and Affective Neuroscience, 16(7), 645–656. https://doi.org/10.1093/scan/nsab029.

Clarke, G., Wright, D.M., 1984. A comparison of analgesia and suppression of oxytocin release by opiates. Br. J. Pharmacol. 83, 799. https://doi.org/10.1111/J.1476-5381.1984.TB16235.X.

Coyle, J.T., Pert, C.B., 1976. Ontogenetic development of [3H]-naloxone binding in rat brain. Neuropharmacology 15, 555–560. https://doi.org/10.1016/0028-3908(76)90107-6.

Dölen, G., Darvishzadeh, A., Huang, K.W., Malenka, R.C., 2013. Social reward requires coordinated activity of nucleus accumbens oxytocin and serotonin. Nature 501, 179–184. https://doi.org/10.1038/nature12518.

Douglas, A.J., Russell, J.A., 2001. Endogenous opioid regulation of oxytocin and ACTH secretion during pregnancy and parturition, in: Progress in Brain Research. Elsevier, pp. 67–82. https://doi.org/10.1016/S0079-6123(01)33006-6.

Fill, M.M.A., Miller, A.M., Wilkinson, R.H., Warren, M.D., Dunn, J.R., Schaffner, W., Jones, T.F., 2018. Educational disabilities among children born with neonatal abstinence syndrome. Pediatrics 142, e20180562. https://doi.org/10.1542/peds.2018-0562.

Gagin, R., Kook, N., Cohen, E., Shavit, Y., 1997. Prenatal morphine enhances morphine-conditioned place preference in adult rats. Pharmacology, Biochemistry, and Behavior, 58(2), 525–528. https://doi.org/10.1016/s0091-3057(97)00281-5.

Hirai, A.H., Ko, J.Y., Owens, P.L., Stocks, C., Patrick, S.W., 2021. Neonatal abstinence syndrome and maternal opioid-related diagnoses in the US, 2010-2017. JAMA - J. Am. Med. Assoc. 325, 146–155. https://doi.org/10.1001/jama.2020.24991.

Hol, T., Niesink, M., Van Ree, J.M., Spruijt, B.M., 1996. Prenatal exposure to morphine affects juvenile play behavior and adult social behavior in rats. Pharmacol. Biochem. Behav. 55, 615–618. https://doi.org/10.1016/S0091-3057(96)00274-2.

Hunt, R. W., Tzioumi, D., Collins, E., Jeffery, H. E., 2008. Adverse neurodevelopmental outcome of infants exposed to opiate in-utero. Early Human Development, 84(1), 29–35. https://doi.org/10.1016/j.earlhumdev.2007.01.013.

Inenaga, K., Imura, H., Yanaihara, N., Yamashita, H., 1990. Kappa-selective opioid receptor agonists leumorphin and dynorphin inhibit supraoptic neurons in rat hypothalamic slice preparations. J. Neuroendocrinol. 2, 389–395. https://doi.org/10.1111/j.1365-2826.1990.tb00423.x.

Jalowiec, J.E., Calcagnetti, D.J., Fanselow, M.S., 1989. Suppression of juvenile social behavior requires antagonism of central opioid systems. Pharmacol. Biochem. Behav. 33, 697–700. https://doi.org/10.1016/0091-3057(89)90409-7.

Johnson, Z.V., Walum, H., Xiao, Y., Riefkohl, P.C., Young, L.J., 2017. Oxytocin receptors modulate a social salience neural network in male prairie voles. Hormones and Behavior 87, 16–24. https://doi.org/10.1016/j.yhbeh.2016.10.009.

Jordan, R., 2003. Social play and autistic spectrum disorders: A perspective on theory, implications and educational approaches. Autism, https://doi.org/10.1177/1362361303007004002.

Keebaugh, A.C., Barrett, C.E., LaPrairie, J.L., Jenkins, J.J., Young, L.J., 2015. RNAi knockdown of oxytocin receptor in the nucleus accumbens inhibits social attachment and parental care in monogamous female prairie voles. Soc Neurosci 10, 561–570. https://doi.org/10.1080/17470919.2015.1040893.

Keebaugh, A.C., Young, L.J., 2011. Increasing oxytocin receptor expression in the nucleus accumbens of pre-pubertal female prairie voles enhances alloparental responsiveness and partner preference formation as adults. Hormones and Behavior 60, 498–504. https://doi.org/10.1016/j.yhbeh.2011.07.018.

Kocherlakota, P., 2014. Neonatal abstinence syndrome. Pediatrics 134, e547–e561. https://doi.org/10.1542/PEDS.2013-3524.

Kuhn, C.M., Windh, R.T., Little, P.J., 1992. Effects of perinatal opiate addiction on neurochemical development of the brain, in: Miller, M.W. (Ed.), Development of the Central Nervous System: Effects of Alcohol and Opiates. Wiley-Liss, New York, pp. 341–361.

Laorden, M.L., Milanés, M. V., Chapleur-Château, M., Burlet, A., 1997. Changes in hypothalamic oxytocin levels during morphine tolerance. Neuropeptides 31, 143–146. https://doi.org/10.1016/S0143-4179(97)90083-4.

Laorden, M.L., Milanés, M. V., Chapleur-Château, M., Burlet, A., 1998. Changes in oxytocin content in rat brain during morphine withdrawal. Neuropeptides 32, 67–71. https://doi.org/10.1016/S0143-4179(98)90019-1.

Li, X., Morrow, D., Witkin, J.M., 2006. Decreases in nestlet shredding of mice by serotonin uptake inhibitors: Comparison with marble burying. Life Sciences 78, 1933–1939. https://doi.org/10.1016/j.lfs.2005.08.002.

Li, J., You, Z., Chen, Z., Song, C., Lu, C., 2001. Chronic morphine treatment inhibits oxytocin release from the supraoptic nucleus slices of rats. Neurosci. Lett. 300, 54–58. https://doi.org/10.1016/S0304-3940(01)01540-3.

Lutz-Bucher, B., Koch, B., 1980. Evidence for a direct inhibitory effect of morphine on the secretion of posterior pituitary hormones. Eur. J. Pharmacol. 66, 375–378. https://doi.org/10.1016/0014-2999(80)90470-7.

Maguire, D.J., Taylor, S., Armstrong, K., Shaffer-Hudkins, E., Germain, A.M., Brooks, S.S., Cline, G.J., Clark, L., 2016. Long-term outcomes of infants with neonatal abstinence syndrome. Neonatal Network 35, 277–286. https://doi.org/10.1891/0730-0832.35.5.277.

Manduca, A., Campolongo, P., Palmery, M., Vanderschuren, L.J.M.J., Cuomo, V., Trezza, V., 2014. Social play behavior, ultrasonic vocalizations and their modulation by morphine and amphetamine in Wistar and Sprague-Dawley rats. Psychopharmacology (Berl). 231, 1661–1673. https://doi.org/10.1007/s00213-013-3337-9.

Meyer-Lindenberg, A., Domes, G., Kirsch, P., Heinrichs, M., 2011. Oxytocin and vasopressin in the human brain: Social neuropeptides for translational medicine. Nat. Rev. Neurosci. https://doi.org/10.1038/nrn3044.

Minakova, E., Mikati, M.O., Madasu, M.K., Conway, S.M., Baldwin, J.W., Swift, R.G., McCullough, K.B., Dougherty, J.D., Maloney, S.E., Al-Hasani, R., 2022. Perinatal oxycodone exposure causes long-term sex-dependent changes in weight trajectory and sensory processing in adult mice. Psychopharmacology (Berl) 239, 3859–3873. https://doi.org/10.1007/s00213-022-06257-8.

Motulsky, H.J., 2021. How to report the methods used for mixed model analysis. https://www.graphpad.com/guides/prism/latest/statistics/stat_how-to-report-the-methods-used.htm (accessed 9 September 2021).

Najam, N., Panksepp J., 1989. Effect of chronic neonatal morphine and naloxone on sensorimotor and social development of young rats. Pharmacology, Biochemistry and Behavior 33:539–544.

Niesink, R.J.M., Van Buren-Van Duinkerken, L., Van Ree, J.M., 1999. Social behavior of juvenile rats after in utero exposure to morphine: Dose-time-effect relationship. Neuropharmacology 38, 1207–1223. https://doi.org/10.1016/S0028-3908(99)00050-7.

Niesink, R.J.M., Van Ree, J.M., 1989. Involvement of opioid and dopaminergic systems in isolation-induced pinning and social grooming of young rats. Neuropharmacology 28, 411–418. https://doi.org/10.1016/0028-3908(89)90038-5.

Niles, J.K., Gudin, J., Radcliff, J., Kaufman, H.W., 2021. The opioid epidemic within the COVID-19 pandemic: Drug testing in 2020. Population Health Management 24, S–43. https://doi.org/10.1089/pop.2020.0230.

Normansell, L., Panksepp, J., 1990. Effects of morphine and naloxone on play-rewarded spatial discrimination in juvenile rats. Dev. Psychobiol. 23, 75–83. https://doi.org/10.1002/dev.420230108.

Panksepp, J., 1981. The ontogeny of play in rats. Dev. Psychobiol. 14, 327–332. https://doi.org/10.1002/dev.420140405.

Parra-Vargas, M., Ramon-Krauel, M., Lerin, C., Jimenez-Chillaron, J. C., 2020. Size does matter: Litter size strongly determines adult metabolism in rodents. Cell Metabolism, 32(3), 334–340. https://doi.org/10.1016/j.cmet.2020.07.014.

Pek, J., Flora, D.B., 2018. Reporting effect sizes in original psychological research: A discussion and tutorial. Psychol. Methods 23, 208–225. https://doi.org/10.1037/met0000126.

Pellis, S. M., Field, E. F., Smith, L. K., & Pellis, V. C. (1997). Multiple differences in the play fighting of male and female rats: Implications for the causes and functions of play. Neuroscience & Biobehavioral Reviews 21(1), 105–120. https://doi.org/10.1016/0149-7634(95)00060-7.

Ramsey, N. F., Niesink, R. J. M., Van Ree, J. M., 1993. Prenatal exposure to morphine enhances cocaine and heroin self-administration in drug-naive rats. Drug and Alcohol Dependence, 33(1), 41–51. https://doi.org/10.1016/0376-8716(93)90032-L.

Reppucci, C.J., Gergely, C.K., Veenema, A.H., 2018. Activation patterns of vasopressinergic and oxytocinergic brain regions following social play exposure in juvenile male and female rats. J. Neuroendocrinol. 30, e12582. https://doi.org/10.1111/ine.12582.

Rigney, N., de Vries, G.J., Petrulis, A., Young, L.J., 2022. Oxytocin, vasopressin, and social behavior: From neural circuits to clinical opportunities. Endocrinology 163. https://doi.org/10.1210/endocr/bqac111.

Sandtorv, L.B., Fevang, S.K.E., Nilsen, S.A., Bøe, T., Gjestad, R., Haugland, S., Elgen, I.B., 2018. Symptoms associated with attention deficit/hyperactivity disorder and autism spectrum disorders in school-aged children prenatally exposed to substances. Subst. Abus. Res. Treat. 12. https://doi.org/10.1177/1178221818765773.

Schwartz, C. L., Christiansen, S., Vinggaard, A. M., Axelstad, M., Hass, U., Svingen, T., 2019. Anogenital distance as a toxicological or clinical marker for fetal androgen action and risk for reproductive disorders. Archives of Toxicology, 93(2), 253–272. https://doi.org/10.1007/s00204-018-2350-5.

Shen, Y.L., Chen, S.T., Chan, T.Y., Hung, T.W., Tao, P.L., Liao, R.M., Chan, M.H., Chen, H.H., 2016. Delayed extinction and stronger drug-primed reinstatement of methamphetamine seeking in rats prenatally exposed to morphine. Neurobiology of Learning and Memory, 128, 56–64. https://doi.org/10.1016/j.nlm.2015.12.002.

Siegel, M.A., Jensen, R.A., 1986. The effects of naloxone and cage size on social play and activity in isolated young rats. Behav. Neural Biol. 45, 155–168. https://doi.org/10.1016/S0163-1047(86)90739-9.

Siegel, M.A., Jensen, R.A., Panksepp, J., 1985. The prolonged effects of naloxone on play behavior and feeding in the rat. Behav. Neural Biol. 44, 509–514. https://doi.org/10.1016/S0163-1047(85)91024-6.

Smith, K., Lipari, R.N., 2017. Women of childbearing age and opioids. The CBHSQ Report: January 17, 2017. Center for Behavioral Health Statistics and Quality, Substance Abuse and Mental Health Services Administration, Rockville, MD. https://www.samhsa.gov/data/sites/default/files/report_2724ZShortReport-2724.html.

Smith, C.J.W., Mogavero, J.N., Tulimieri, M.T., Veenema, A.H., 2017. Involvement of the oxytocin system in the nucleus accumbens in the regulation of juvenile social novelty-seeking behavior. Hormones and Behavior 93, 94–98. https://doi.org/10.1016/j.yhbeh.2017.05.005.

Torabi, M., Pooriamehr, A., Bigdeli, I., & Miladi-Gorji, H., 2017. Maternal swimming exercise during pregnancy attenuates anxiety/depressive-like behaviors and voluntary morphine consumption in the pubertal male and female rat offspring born from morphine dependent mothers. Neuroscience Letters, 659, 110–114. https://doi.org/10.1016/j.neulet.2017.08.074.

Trezza, V., Damsteegt, R., Marijke Achterberg, E.J., Vanderschuren, L.J.M.J., 2011. Nucleus accumbens μ-opioid receptors mediate social reward. J. Neurosci. 31, 6362–6370. https://doi.org/10.1523/JNEUROSCI.5492-10.2011.

Van de Heijning, B.J.M., Van den Herik, I.K., Van Wimersma Greidanus, T.B., 1991. The opioid receptor subtypes μ and κ, but not δ, are involved in the control of the vasopressin and oxytocin release in the rat. Eur. J. Pharmacol. 209, 199–206. https://doi.org/10.1016/0014-2999(91)90170-U.

Van den Berg, C.L., Pijlman, F.T., Koning, H.A., Diergaarde, L., Van Ree, J.M., Spruijt, B.M., 1999a. Isolation changes the incentive value of sucrose and social behaviour in juvenile and adult rats. Behav Brain Res 106:133–142. https://doi.org/10.1016/s0166-4328(99)00099-6.

Van Den Berg, C.L., Van Ree, J.M., Spruijt, B.M., 1999b. Sequential analysis of juvenile isolation-induced decreased social behavior in the adult rat. Physiol Behav 67:483–488. https://doi.org/10.1016/S0031-9384(99)00062-1.

Van Den Berg, C.L., Van Ree, J.M., Spruijt, B.M., 2000. Morphine attenuates the effects of juvenile isolation in rats. Neuropharmacology 39, 969–976. https://doi.org/10.1016/S0028-3908(99)00216-6.

Vanderschuren, L.J.M.J., Niesink, R.J.M., Spruijt, B.M., Van Ree, J.M., 1995a. μ- and κ-opioid receptor-mediated opioid effects on social play in juvenile rats. Eur. J. Pharmacol. 276, 257–266. https://doi.org/10.1016/0014-2999(95)00040-R.

Vanderschuren, L.J.M.J., Spruijt, B.M., Hol, T., Niesink, R.J.M., Van Ree, J.M., 1995b. Sequential analysis of social play behavior in juvenile rats: effects of morphine. Behav. Brain Res. 72, 89–95. https://doi.org/10.1016/0166-4328(96)00060-5.

Vanderschuren, L.J.M.J., Spruijt, B.M., Van Ree, J.M., Niesink, R.J.M., 1995c. Effects of morphine on different aspects of social play in juvenile rats. Psychopharmacology (Berl). 117, 225–231. https://doi.org/10.1007/BF02245191.

Vanderschuren, L.J.M.J., Stein, E.A., Wiegant, V.M., Van Ree, J.M., 1995d. Social play alters regional brain opioid receptor binding in juvenile rats. Brain Res. 680, 148–156. https://doi.org/10.1016/0006-8993(95)00256-P.

Van Ree, J.M., Niesink, R.J.M., 1983. Low doses of β-endorphin increase social contacts of rats tested in dyadic encounters. Life Sci. 33, 611–614. https://doi.org/10.1016/0024-3205(83)90577-5.

Veenema, A.H., 2012. Toward understanding how early-life social experiences alter oxytocin- and vasopressin-regulated social behaviors. Horm Behav 61, 304–312. https://doi.org/10.1016/j.yhbeh.2011.12.002.

Veenema, A. H., Bredewold, R., De Vries, G. J., 2013. Sex-specific modulation of juvenile social play by vasopressin. Psychoneuroendocrinology, 38(11), 2554–2561. https://doi.org/10.1016/j.psyneuen.2013.06.002.

Wilkinson, L., 1999. Statistical methods in psychology journals: Guidelines and explanations. Am. Psychol. 54, 594–604. https://doi.org/10.1037/0003-066X.54.8.594.

Wu, L.Y., Chen, J.F., Tao, P.L., Huang, E.Y.K., 2009. Attenuation by dextromethorphan on the higher liability to morphine-induced reward, caused by prenatal exposure of morphine in rat offspring. Journal of Biomedical Science, 16(1), 106. https://doi.org/10.1186/1423-0127-16-106.

Yeoh, S.L., Eastwood, J., Wright, I.M., Morton, R., Melhuish, E., Ward, M., Oei, J.L., 2019. Cognitive and motor outcomes of children with prenatal opioid exposure: A systematic review and meta-analysis. JAMA Network Open 2, e197025. https://doi.org/10.1001/jamanetworkopen.2019.7025.

You, Z.D., Li, J.H., Song, C.Y., Wang, C.H., Lu, C.L., 2000. Chronic morphine treatment inhibits oxytocin synthesis in rats. Neuroreport 11, 3113–3116. https://doi.org/10.1097/00001756-200009280-00015.

Zhang, R., Zhang, H.F., Han, J.S., Han, S.P., 2017. Genes related to oxytocin and arginine-vasopressin pathways: Associations with autism spectrum disorders. Neurosci. Bull. https://doi.org/10.1007/s12264-017-0120-7.

Zhao, B.G., Chapman, C., Bicknell, R.J., 1988. Functional κ-opioid receptors on oxytocin and vasopressin nerve terminals isolated from the rat neurohypophysis. Brain Res. 462, 62–66. https://doi.org/10.1016/0006-8993(88)90585-9.

